## Supplementary material for "Anti-CSF-1R therapy with combined immuno- chemotherapy coordinate an adaptive immune response to eliminate macrophage enriched Triple Negative Breast Cancers"

**Supplementary Figure 1. CSF-1R inhibition coupled with CTX leads to a durable response in primary macrophage enriched TNBC pre-clinical GEMMs.** (A) Schematic illustration of IMC protocol, including isotope labeling, laser ablation and the antibody body panel that includes marker that identify tumor, proliferation, checkpoint, lymphocytes and monocytes. Analysis including single cell level profiling and TSNE were utilized to study the TIME. (B) Heatmap depicting 20 different clusters (cluster_id) from individual protein markers with the respective number of cells (n_cells) detected. (C) Double IF staining of macrophages (F4/80) and neutrophils (S100A8) from claudin-low (T12, 2151R and T11), basal-like (2336R) and luminal-like (2208L) GEMMs. (D) IHC staining of macrophages (F4/80) following IgG control or SNDX-ms6352 treatment in T12 model. (E) IHC quantification of macrophages (F4/80) following IgG control or SNDX-ms6352 treatment in T12 model. Statistical analysis performed using 2-tailed, unpaired Student's t test. ($n$ = 8). (F) IF staining of macrophages (F4/80) and apoptosis (Cleaved Caspase 3) following IgG control or SNDX-ms6352 treatment in T12 model ($n$ = *3* mice per treatment). (G) Tumor volume following IgG control or SNDX-ms6352 treatment in T12 model. ($n$ = *8* mice per treatment). (H) Immunoblots of TAMs isolated from T12 and 2151R TAMs treated with increasing dose of IgG control or SNDX-ms6352. (I) IF staining of cytoskeleton (phalloidin) and CSF-1R marker following IgG control or SNDX-ms6352 treatment in isolated TAMs *in vitro* ($n$ = *3*). (J) Tumor volume of 2151R contralateral tumor cell rechallenged mice post CTX or combination treatment. Statistical analysis performed using Two-way ANOVA, ($n$ = 2-5 mice per treatment). (K) H&E and IF staining of B cell (CD20) and T cell (CD8) following contralateral tumor cell rechallenged mice post combination treatment in the T12 model ($n$ = 3 mice). Data shown as mean ±SEM. For all panels *$P$ < 0.05; **$P$ < 0.01; *** $P$ < 0.001; ****$P$ < 0.0001.

**Supplementary Figure 2. CSF-1R inhibition coupled with CTX leads to a durable response in primary macrophage enriched TNBC pre-clinical GEMMs.** (A) scRNA-seq transcriptomic expression of TAM markers from T12 and 2151R models following IgG control, SNDX-ms6352, CTX and the combination. (B) UMAP clustering from scRNA-seq of *Fabp5* expression levels in various residual TAMs following IgG control, SNDXms6352, CTX and the combination in the T12 model. (C) UMAP clustering of different TAM populations identified via scRNA-seq within the 2151R and T12 models. (D) Flow cytometry of F4/80$^+$ macrophages within CD45$^+$ immune cells in the T12 model following IgG control, SNDX-ms6352, CTX and the combination. Statistical analysis performed using Two-way ANOVA, ($n$ = 5 mice per treatment). (E) Cytokine array profiling from the plasma of SNDX-ms6352, CTX and combination treated mice. Median expression levels compared to IgG controls (fold change), ($n$ = 3 mice per treatment). (F) Cytokine array profiling from the tumor lysate of T12 IgG control, SNDX-ms6352, CTX and combination treated mice. Median expression levels compared to IgG controls (fold change) ($n$ = 3 mice per treatment). (G) Tumor volume of T12 tumors transplanted into immunodeficient athymic nude mice and treated with IgG control, SNDX-ms6352, CTX or combination. ($n$ = 5-7 mice per treatment). (H) Illustration depicting cytokine response observed in plasma and tumor homogenates following each treatment in mice bearing T12 primary tumors. Data shown as mean ±SEM. For all panels *$P$ < 0.05; **$P$ < 0.01; *** $P$ < 0.001; ****$P$ < 0.0001.

**Supplementary Figure 3. Combination treatment leads to ITH driven PD-1/PD-L1 expression in metastatic lung metastases.** (A) Whole lung tissue H&E scans and IF staining of BrdU$^+$ tumor cells post 12-day T12 tumor cell TV injection, (*n = 3* independent mice*)*. (B) IHC staining of macrophage (F4/80) from whole lung tissues scans following IgG control or SNDX-ms6352, *(n = 3* independent mice*)*. (C) 20x magnification IHC staining of macrophage (F4/80) following IgG control or SNDX-ms6352. (D) Cumulative and individual marker (Vimentin, Pan-CK, CD8a, CD44, PD-1, FoxP3, F4/80, S100A8/9, PD-L1 and Ki67) IMC analysis from multiple IgG (control 1 and control 2) treated lung macro-metastases. (E) Bar plots from IMC analysis of IgG control lung MaMets and combination treated lung MiMets looking for neutrophil, PanCK$^+$, PD-1$^+$, PD-L1$^+$ and tumor cell populations. (F) ITH is observed in combination treated MiMet1 and MiMet2 compared to MaMet1 IgG controls following IMC analysis (Vimentin, B220, S100A8, Ki67 and PD-L1). (G) IMC quantification of EMT (Vimentin), proliferation (Ki67) and exhaustion markers (PD-L1) in MiMet1.1, MiMet1.2 and IgG Control. Statistical analysis performed using two-way ANOVA with multiple comparisons test, (*n* = 3 mice). (H) H&E and IHC staining of macrophages (F4/80) and T cells (CD8) in recurrent lung metastases. (*n = 3* mice) Data shown as mean ±SEM. For all panels **P* < 0.05; ***P* < 0.01; *** *P* < 0.001; *****P* < 0.0001.

**Supplementary Figure 4. Addition of anti-PD1 reshapes the landscape of the metastatic TIME leading to the proximity of B-, T-cells and neutrophils and treatment response in established lung metastases.** (A) Lung wholemounts from T12 IgG control, SNDX-ms6352, CTX and double or triple combination treated lungs, (*n = 5* mice per treatment). (B) Heatmap depicting 20 different clusters from individual protein markers (cluster_id) with the representative number of cells (n_cells) detected. (C) TSNE visualization of macrophage, neutrophil and tumor cells following IgG control, SNDX-ms6352, CTX and double or triple combination treated T12 treated lung metastases. (D) Median expression levels of PanCK, CD44, F4/80, S100A8/9, Ki67, CD4, CD8a and Vimentin following IgG control, SNDX-ms6352, CTX and double or triple combination treated T12 treated lung metastases. *P*-values were generated using diffcyt function (type of analysis, DS; method for DS testing, diffcyt-DS-limma). *P*-values were then adjusted using the Benjamini-Hochberg method, (*n* = 3 ROI). (E) IMC analysis of T cells (CD4 and CD8a) and neutrophils (S100A9) following IgG control, SNDX-ms6352, CTX and double or triple combination treated T12 treated lung metastases, (*n* = 3 ROI). (F) Unsupervised clustering of single cells segmented from IMC data from IgG control T12 lung metastases resulted in 20 clusters. (G) Expression levels of single cells segmented from IMC data from IgG control, SNDX-ms6352, CTX, CTX+SNDX-ms6352 and CTX+SNDX-ms6352+aPD-1 in T12 lung metastases from 20 clusters.

**Supplementary Figure 5. Triple combination leads to long-term durable response of established lung metastases.** (A) Double IF staining of CSF-1R, BrdU, CD20, CD8 and DAPI in T12 mice with established lung metastases following IgG control, SNDX-ms6352, CTX, CTX+SNDX-ms6352 or CTX+SNDX-ms6352+aPD-1, (*n = 5* mice per treatment). (B) IHC staining of macrophages (F4/80), neutrophils (S100A8) and T cells

(CD4 and CD8) in T12 mice with established lung metastases following IgG control, SNDX-ms6352, CTX, CTX+SNDX-ms6352 or CTX+SNDX-ms6352+aPD-1, (*n = 5* mice per treatment). (C) IHC quantification of macrophages (F4/80), neutrophils (S100A8) and T cells (CD4 and CD8) from T12 mice with established lung metastases following IgG control, SNDX-ms6352, CTX, CTX+SNDX-ms6352 or CTX+SNDX-ms6352+aPD-1. Statistical analysis was performed using two-way ANOVA multiple comparisons test, ($n$ = 5 mice per treatment). Data shown as mean ±SEM. For all panels *$P$ < 0.05; **$P$ < 0.01; *** $P$ < 0.001; ****$P$ < 0.0001.

**Supplementary Figure 6. Established T12 liver metastases present with an immunologically cold TIME.** (A) Liver wholemounts from T12 IgG control, SNDX-ms6352, CTX and double or triple combination treated lungs, (*n = 3-5* mice per treatment). (B) Heatmap depicting 20 different clusters from individual protein markers with the representative number of cells. (C) Unsupervised clustering of single cells segmented from IMC data from IgG control T12 liver metastases resulted in 20 clusters. (D) Expression levels of single cells segmented from IMC data from IgG control, SNDX-ms6352, CTX, CTX+SNDX-ms6352 and CTX+SNDX-ms6352+aPD-1 in T12 lung metastases from 20 clusters. (E) Median distance analysis of immune cells from tumor cells post IgG, SNDX-ms6352, CTX, CTX+SNDX-ms6352 and CTX+SNDX-ms6352+aPD1 treated T12 GEMMs at day 7, distance analysis is representative from IMC analysis of individual channels CD8, CD4, S100A8/9 and F4/80. Significance was determined using Two-way ANOVA and Tukey's multiple comparisons tests ($n$ = 3 ROI). (F) Cumulative and individual marker (Vimentin, CD44, Ki67, Ly6G and S100A8/9) IMC analysis from multiple triple combination treated liver metastases. ($n$ = 3 ROI). (G) IMC analysis and quantification of macrophages (F4/80) within the tumor bed and stroma of liver metastasis post SNDX-ms6352. Statistical analysis performed using paired t-test ($n$ = 3 ROI). Data shown as mean ±SEM. For all panels *$P$ < 0.05; **$P$ < 0.01; *** $P$ < 0.001; ****$P$ < 0.0001.

**Supplementary Figure 7. An anti-PD-1 maintenance dose is necessary to generates long-term anti-tumor immunity and eradicate established liver metastases.** (A) Double IF staining of CSF-1R, BrdU, CD20, CD8 and DAPI in T12 mice with established liver metastases following IgG control, SNDX-ms6352, CTX, CTX+SNDX-ms6352 (*n = 3* mice per treatment). (B) IHC staining of macrophages (F4/80), neutrophils (S100A8) and T cells (CD4 and CD8) in T12 mice with established liver metastases following IgG control, SNDX-ms6352, CTX, CTX+SNDX-ms6352 (*n = 3* mice per treatment. (C) IHC quantification of macrophages (F4/80), neutrophils (S100A8) and T cells (CD4 and CD8) from T12 mice with established lung metastases following IgG control, SNDX-ms6352, CTX, CTX+SNDX-ms6352. Statistical analysis was performed using two-way ANOVA multiple comparisons test, (*n = 3* mice per treatment). (D) Schematic representation of tumor cell rechallenge experiment. Post PV injection and treatment cessation at day 28, CR mice were rechallenged with 1,000 freshly dissociated tumor cells in the mammary gland. (E) Individual spider plots depicting tumor volume following IgG control, anti-PD-1, IgG control + T cells or anti-PD-1 + T cells from T12 tumor bearing mice following ACT. (F) IHC staining of CD8 T cells in spleen tissues following ACT (*n = 3* mice per treatment). (G) IHC staining of CD8 T cells in T12 primary tumors following ACT (*n = 3* mice per treatment). Data shown as mean ±SEM. For all panels **P* < 0.05; ***P* < 0.01; *** *P* < 0.001; *****P* < 0.0001.

**Supplementary Figure 8. Patients display elevated immunosuppressive macrophage signatures in lung and liver metastatic sites.** (A) Representative heatmap of various immunological and claudin-low/p53 signatures between primary and metastatic tumor in multiple breast cancer subtypes (*n* = 44 primary versus *n* = 8 lung metastatic tumors and *n* = 18 liver metastatic tumors). (B) Volcano plot depicting the most up- and downregulated genes in metastatic samples comparing lung vs primary, liver vs breast and liver vs lung. For all multiple comparisons, p-values were adjusted using the Benjamini-Hochberg method. Gene set enrichment analyses were performed using the fgsea package. Gene sets with adjusted p-values < 0.05 were considered significant. Differential gene expression /expression signature analyses were performed using the limma package. Patient IDs were treated as a corresponding variable.

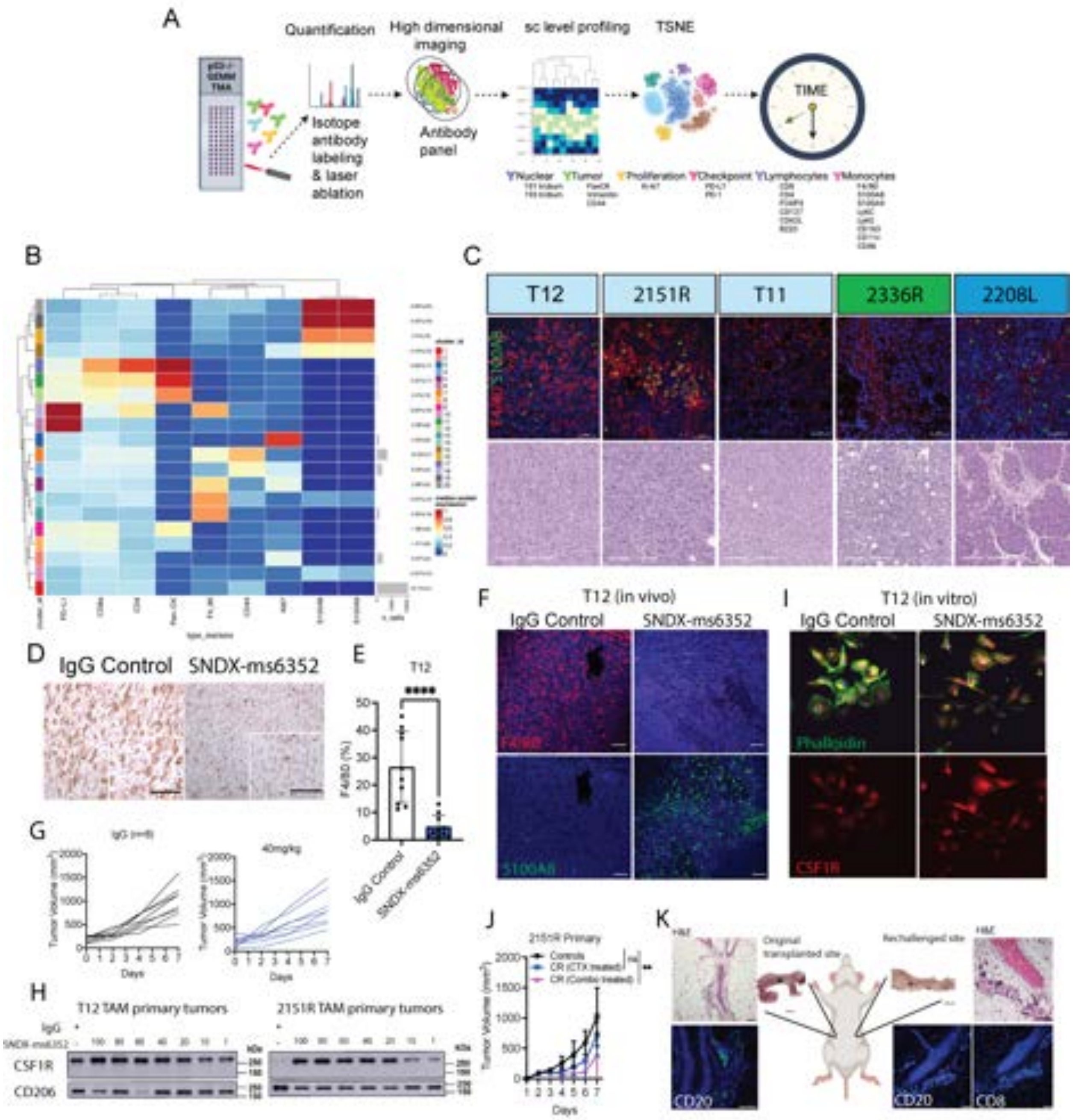
A
Quantification
High dimensional imaging
sc level profiling
TSNE
Isotope antibody labeling & laser ablation
Antibody panel
TIME
Nuclear
Tumor
Proliferation
Checkpoint
Lymphocytes
Monocytes
B
C
T12
2151R
T11
2336R
2208L
F4/80
S100A8
D
IgG Control
SNDX-ms6352
E
T12
****
F4/80 (%)
IgG Control
SNDX-ms6352
F
T12 (in vivo)
IgG Control
SNDX-ms6352
F4/80
S100A8
I
T12 (in vitro)
IgG Control
SNDX-ms6352
Phalloidin
CSF1R
G
Tumor Volume (mm³)
40mg/kg
Days
H
T12 TAM primary tumors
2151R TAM primary tumors
IgG
CSF1R
CD206
kDa
J
2151R Primary
Controls
Tumor Volume (mm³)
Days
K
Original transplanted site
Rechallenged site
H&E
CD20
CD8

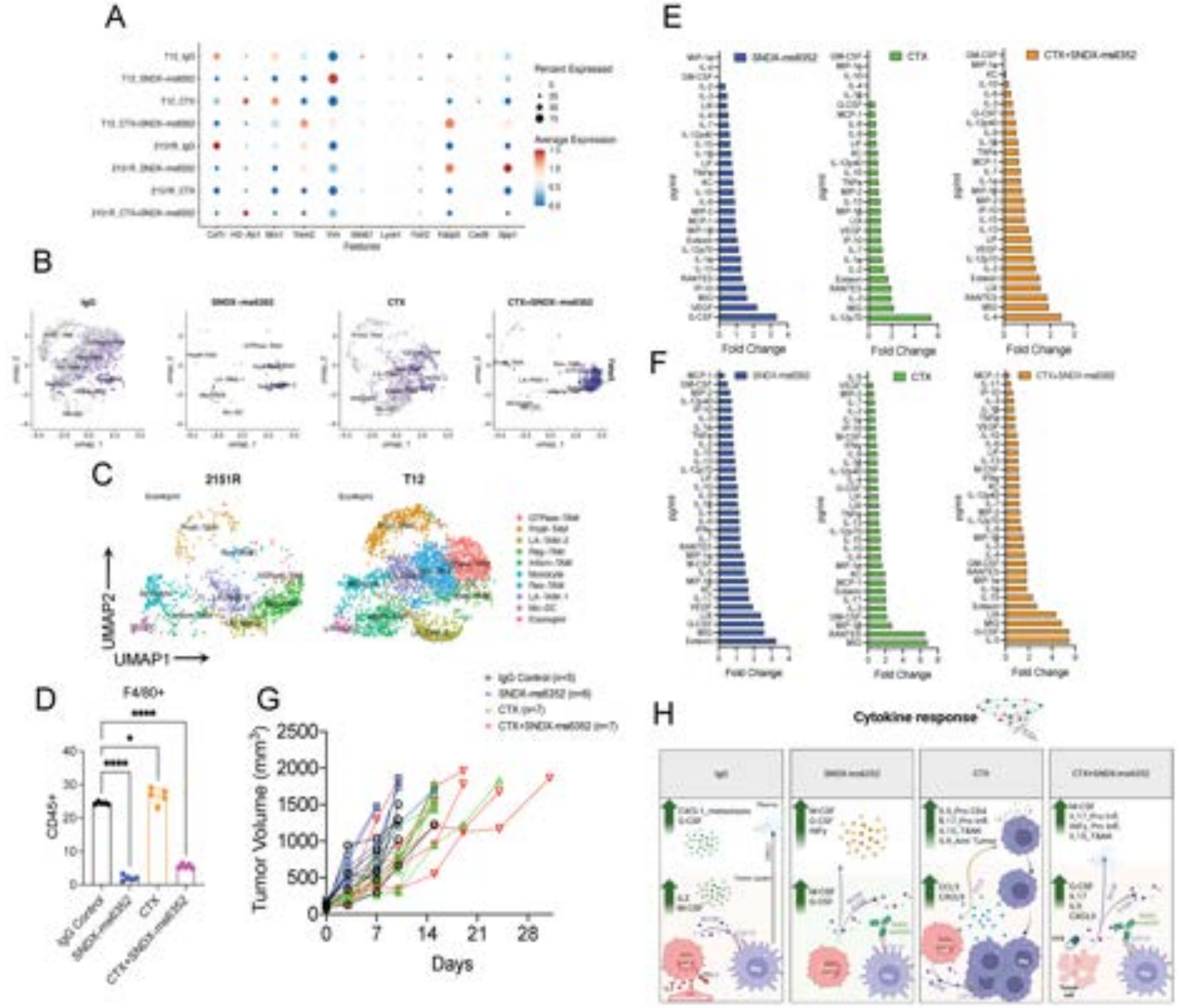
A
B
C
2151R
T12
UMAP2
UMAP1
D
F4/80+
CD45+
IgG Control
SNDX-ms6352
CTX
CTX+SNDX-ms6352
G
IgG Control (n=5)
SNDX-ms6352 (n=6)
CTX (n=7)
CTX+SNDX-ms6352 (n=7)
Tumor Volume (mm3)
Days
E
F
Fold Change
H
Cytokine response

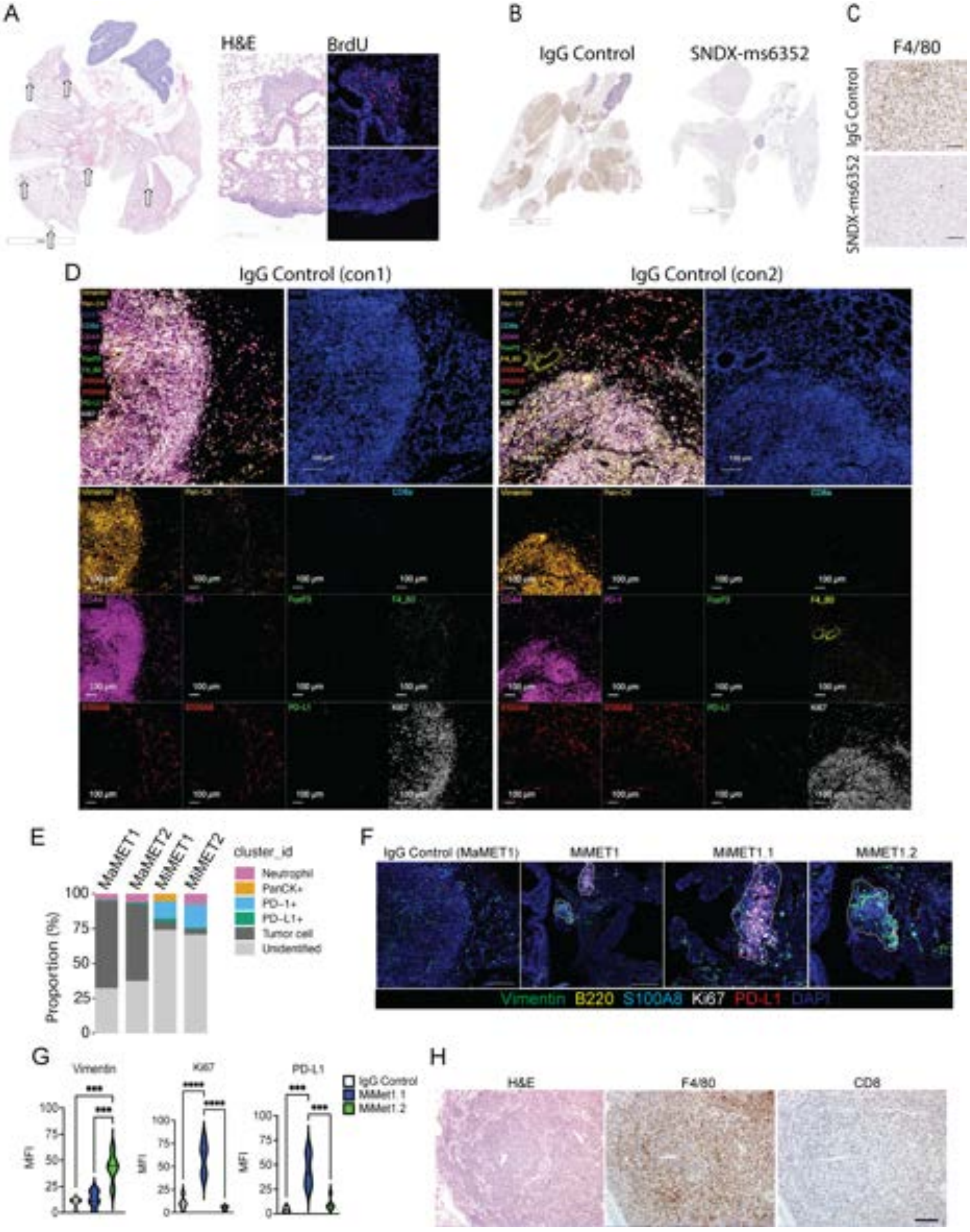

A
H&E
BrdU
B
IgG Control
SNDX-ms6352
C
F4/80
IgG Control
SNDX-ms6352
D
IgG Control (con1)
IgG Control (con2)
E
MaMET1
MaMET2
MiMET1
MiMET2
cluster_id
Neutrophil
PanCK+
PD-1+
PD-L1+
Tumor cell
Unidentified
Proportion (%)
F
IgG Control (MaMET1)
MiMET1
MiMET1.1
MiMET1.2
Vimentin
B220
S100A8
Ki67
PD-L1
DAPI
G
Vimentin
Ki67
PD-L1
IgG Control
MiMet1.1
MiMet1.2
MFI
H
H&E
F4/80
CD8

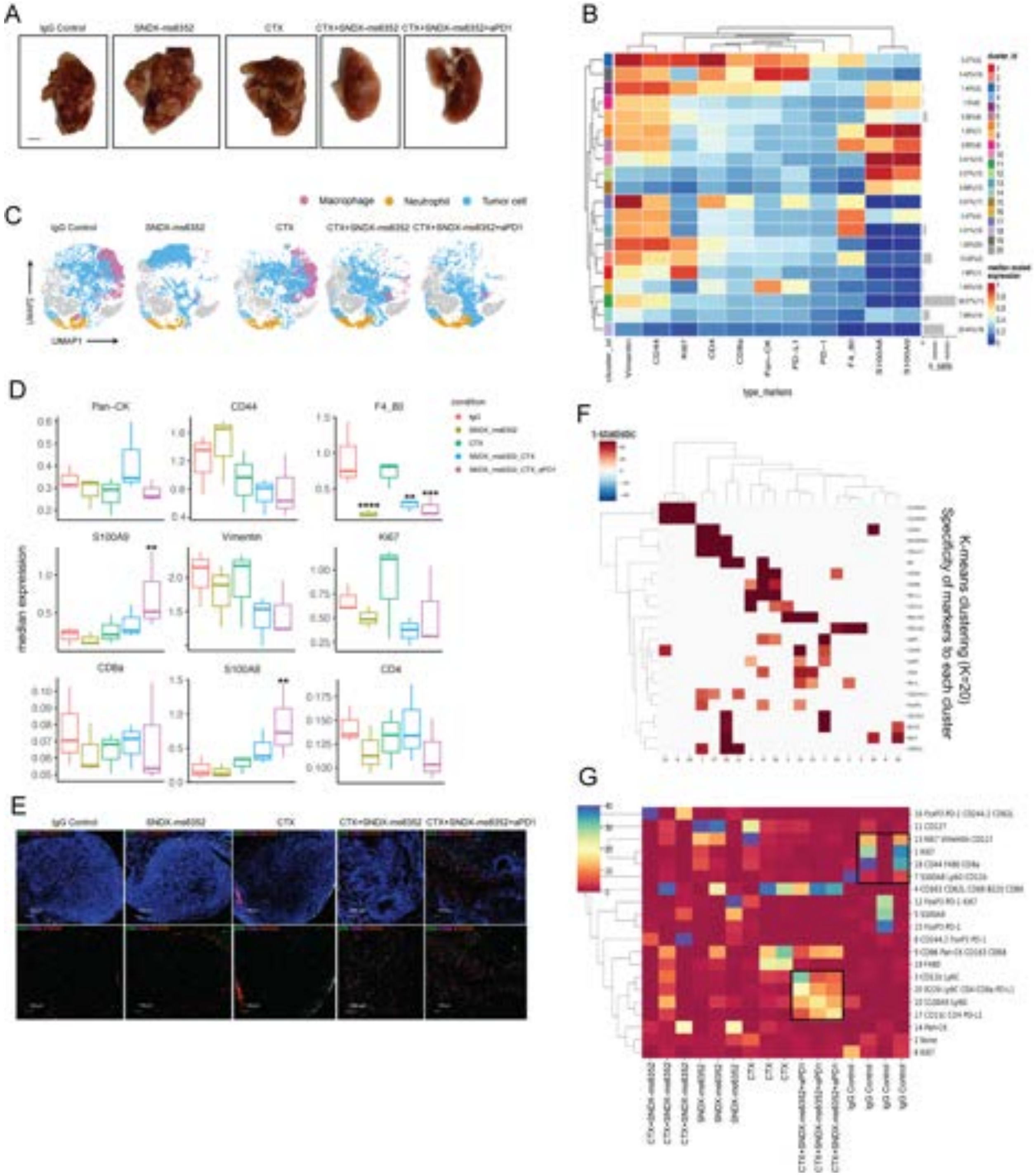

A
IgG Control
SNDX-ms6352
CTX
CTX+SNDX-ms6352
CTX+SNDX-ms6352+aPD1
B
C
Macrophage
Neutrophil
Tumor cell
UMAP1
D
Pan-CK
CD44
F4_80
S100A9
Vimentin
Ki67
CD8a
S100A8
CD4
median expression
E
F
G
K-means clustering (K=20)
Specificity of markers to each cluster

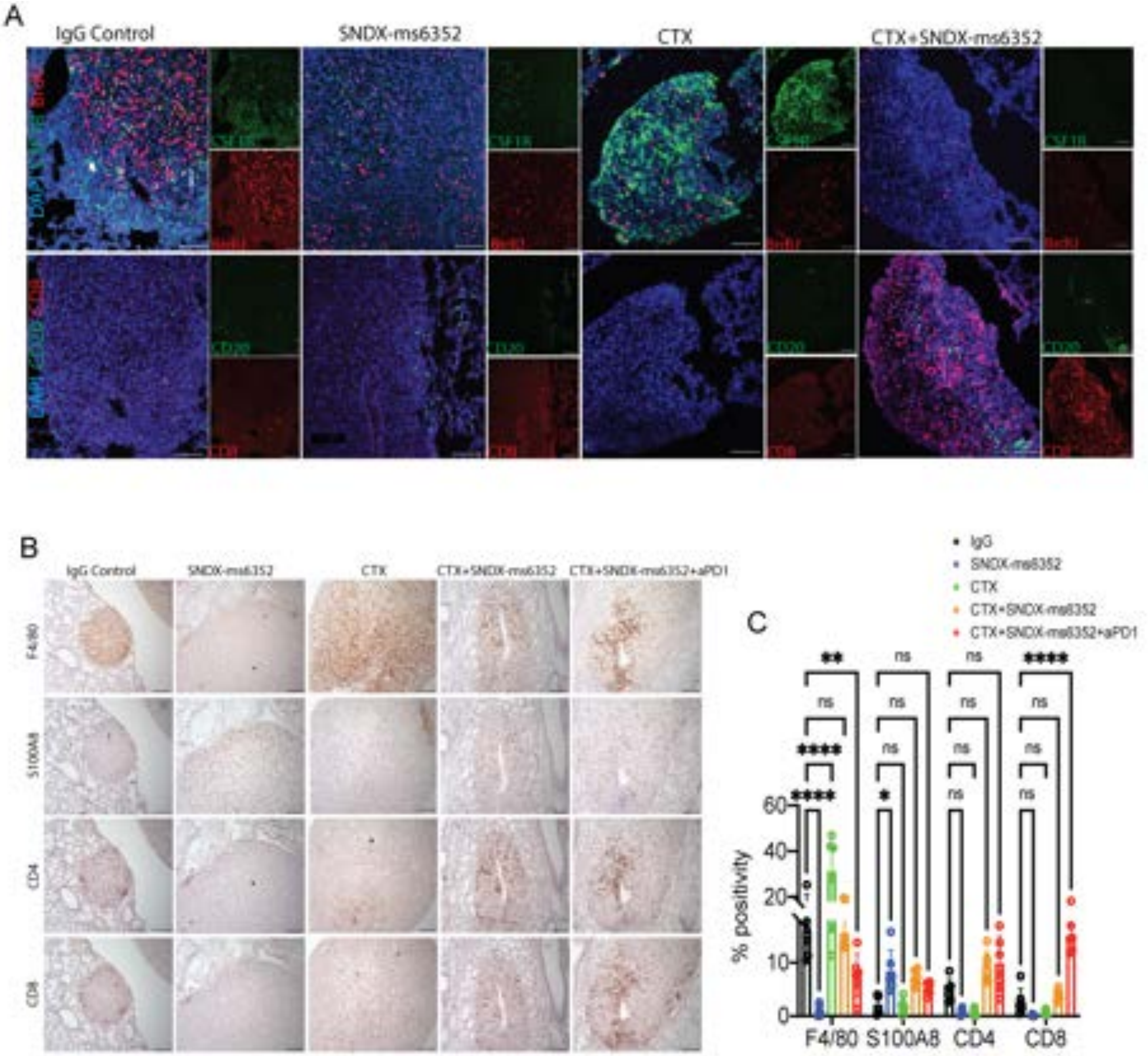
A
IgG Control
SNDX-ms6352
CTX
CTX+SNDX-ms6352
CD20
CD8
B
IgG Control
SNDX-ms6352
CTX
CTX+SNDX-ms6352
CTX+SNDX-ms6352+aPD1
F4/80
S100A8
CD4
CD8
C
IgG
SNDX-ms6352
CTX
CTX+SNDX-ms6352
CTX+SNDX-ms6352+aPD1
% positivity
F4/80
S100A8
CD4
CD8

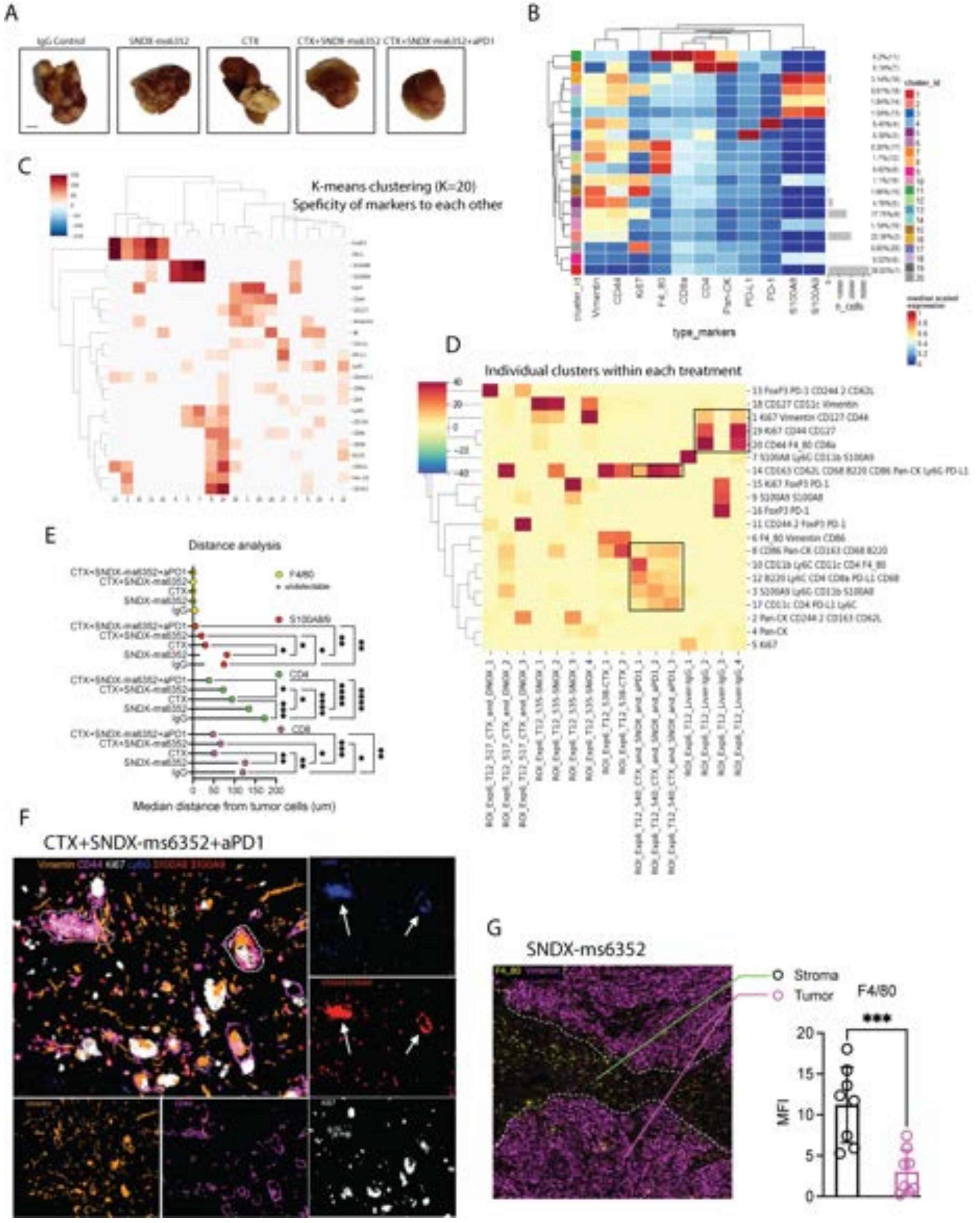
A
IgG Control
SNDX-ms6352
CTX
CTX+SNDX-ms6352
CTX+SNDX-ms6352+aPD1
B
cluster_id
Vimentin
CD44
Ki67
F4_80
CD8a
CD4
Pan-CK
PD-L1
PD-1
S100A8
S100A9
n_cells
type_markers
C
K-means clustering (K=20)
Speficity of markers to each other
D
Individual clusters within each treatment
E
Distance analysis
F4/80
S100A8/9
CD4
CD8
CTX+SNDX-ms6352+aPD1
CTX+SNDX-ms6352
CTX
SNDX-ms6352
IgG
0
50
100
150
200
Median distance from tumor cells (um)
F
CTX+SNDX-ms6352+aPD1
G
SNDX-ms6352
Stroma
Tumor
F4/80
***
MFI
0
5
10
15
20

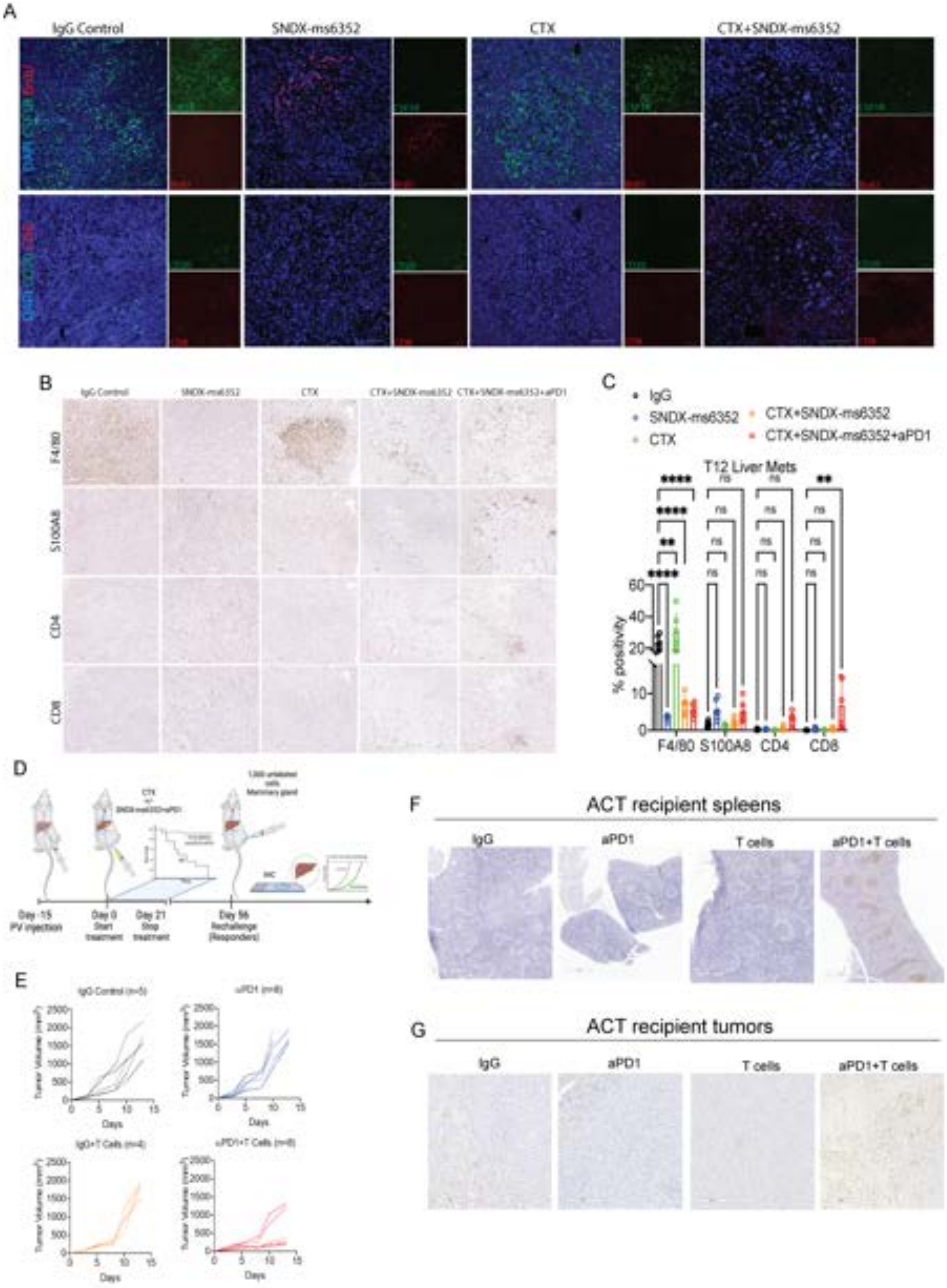
A
IgG Control
SNDX-ms6352
CTX
CTX+SNDX-ms6352
B
F4/80
S100A8
CD4
CD8
C
IgG
SNDX-ms6352
CTX
CTX+SNDX-ms6352
CTX+SNDX-ms6352+aPD1
T12 Liver Mets
% positivity
F4/80
S100A8
CD4
CD8
D
Day -15 PV injection
Day 0 Start treatment
Day 21 Stop treatment
Day 56 Rechallenge (Responders)
E
Tumor Volume (mm³)
Days
F
ACT recipient spleens
IgG
aPD1
T cells
aPD1+T cells
G
ACT recipient tumors
IgG
aPD1
T cells
aPD1+T cells

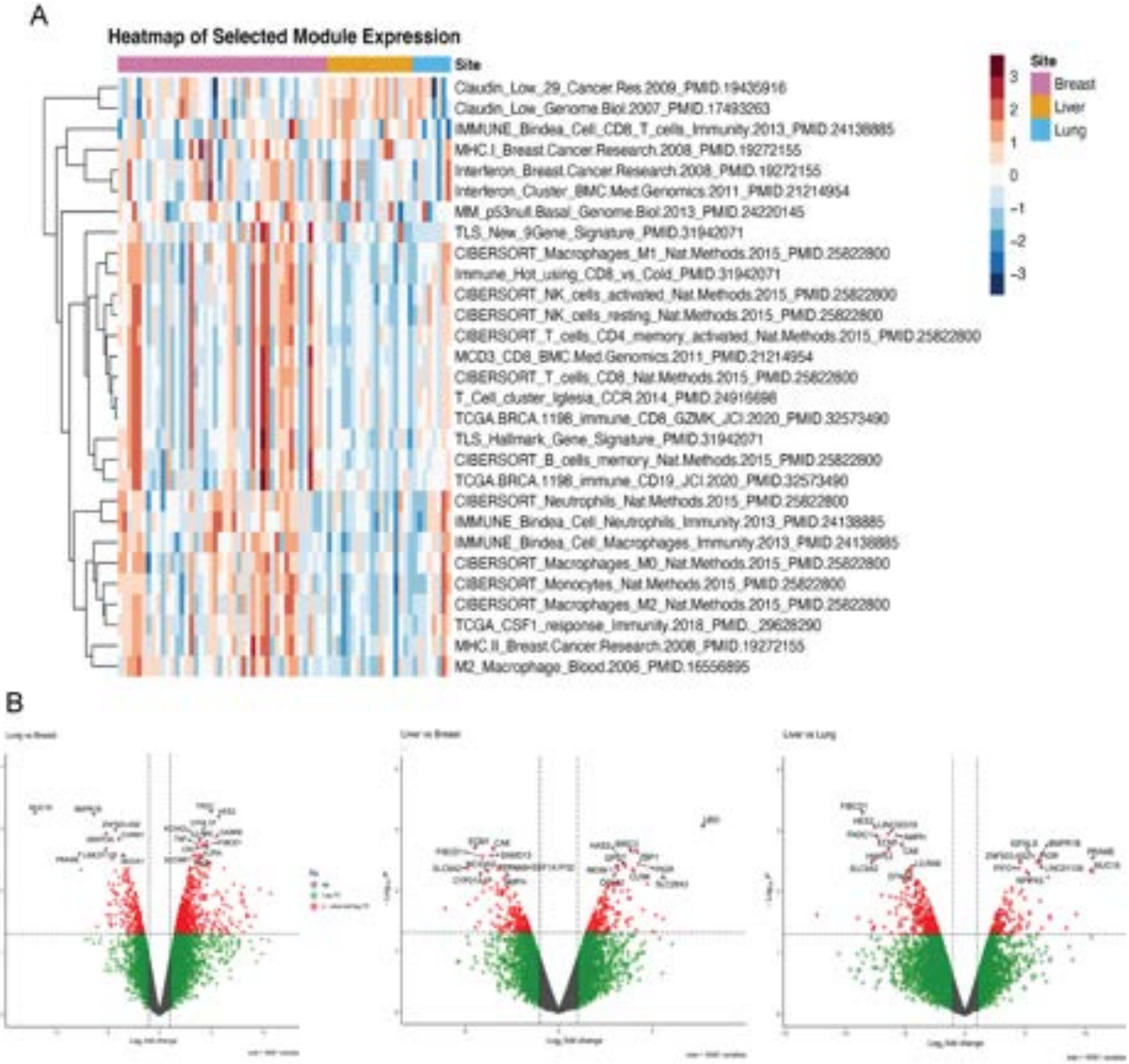
A
Heatmap of Selected Module Expression
Site
Claudin_Low_29_Cancer.Res.2009_PMID.19435916
Claudin_Low_Genome.Biol.2007_PMID.17493263
IMMUNE_Bindea_Cell_CD8_T_cells_Immunity.2013_PMID.24138885
MHC.I_Breast.Cancer.Research.2008_PMID.19272155
Interferon_Breast.Cancer.Research.2008_PMID.19272155
Interferon_Cluster_BMC.Med.Genomics.2011_PMID.21214954
MM_p53null.Basal_Genome.Biol.2013_PMID.24220145
TLS_New_9Gene_Signature_PMID.31942071
CIBERSORT_Macrophages_M1_Nat.Methods.2015_PMID.25822800
Immune_Hot_using_CD8_vs_Cold_PMID.31942071
CIBERSORT_NK_cells_activated_Nat.Methods.2015_PMID.25822800
CIBERSORT_NK_cells_resting_Nat.Methods.2015_PMID.25822800
CIBERSORT_T_cells_CD4_memory_activated_Nat.Methods.2015_PMID.25822800
MCD3_CD8_BMC.Med.Genomics.2011_PMID.21214954
CIBERSORT_T_cells_CD8_Nat.Methods.2015_PMID.25822800
T_Cell_cluster_Iglesia_CCR.2014_PMID.24916698
TCGA.BRCA.1198_immune_CD8_GZMK_JCI.2020_PMID.32573490
TLS_Hallmark_Gene_Signature_PMID.31942071
CIBERSORT_B_cells_memory_Nat.Methods.2015_PMID.25822800
TCGA.BRCA.1198_immune_CD19_JCI.2020_PMID.32573490
CIBERSORT_Neutrophils_Nat.Methods.2015_PMID.25822800
IMMUNE_Bindea_Cell_Neutrophils_Immunity.2013_PMID.24138885
IMMUNE_Bindea_Cell_Macrophages_Immunity.2013_PMID.24138885
CIBERSORT_Macrophages_M0_Nat.Methods.2015_PMID.25822800
CIBERSORT_Monocytes_Nat.Methods.2015_PMID.25822800
CIBERSORT_Macrophages_M2_Nat.Methods.2015_PMID.25822800
TCGA_CSF1_response_Immunity.2018_PMID._29628290
MHC.II_Breast.Cancer.Research.2008_PMID.19272155
M2_Macrophage_Blood.2006_PMID.16556895
3
2
1
0
−1
−2
−3
Site
Breast
Liver
Lung
B
